# Expression of terminal deoxynucleotidyl transferase (TdT) identifies lymphoid-primed progenitors in human bone marrow

**DOI:** 10.1101/2022.10.30.514380

**Authors:** YeEun Kim, Ariel A. Calderon, Patricia Favaro, David R. Glass, Albert G. Tsai, Luciene Borges, William J. Greenleaf, Sean C. Bendall

## Abstract

Lymphoid specification in human hematopoietic progenitors is not fully understood. To better associate lymphoid identity with protein-level cell features, we conducted a highly multiplexed single-cell proteomic screen on human bone marrow progenitors. This screen identified terminal deoxynucleotidyl transferase (TdT), a specialized DNA polymerase intrinsic to VDJ recombination, broadly expressed within CD34+ progenitors prior to B/T cell emergence. While these TdT+ cells coincided with granulocyte-monocyte progenitor (GMP) immunophenotype, their accessible chromatin regions showed enrichment for lymphoid-associated transcription factor (TF) motifs. TdT expression on GMPs was inversely related to the SLAM family member CD84. Prospective isolation of CD84^lo^ GMPs demonstrated robust lymphoid potential *ex vivo*, while still retaining significant myeloid differentiation capacity, akin to LMPPs. This multi-omic study identifies previously unappreciated lymphoid-primed progenitors, redefining the lympho-myeloid axis in human hematopoiesis.

## Introduction

Our understanding of hematopoiesis has changed dramatically in the last decade. Traditionally, hematopoiesis has been portrayed as a hierarchical system in which hematopoietic stem cells (HSCs) differentiate into oligo-potential and uni-potential progenitors in a stepwise manner. And each progenitor population with a distinct differentiation potential was identified by their expression of specific cell surface proteins, also known as ‘surface markers’ (Table 1). However, single-cell techniques revealed the continuous transcriptomic ^1^, epigenetic ^2,3^, and proteomic ^4^ landscapes of human hematopoietic stem and progenitor cells (HSPCs). In parallel, single-cell level differentiation studies demonstrated that cells within the same ‘population’ based on the surface markers exhibit heterogeneous differentiation potentials ^5,6^. Thus, the disagreement between the ideal concept of progenitor cell types with specific differentiation potentials and the empirical cell types identified by their surface immunophenotypes has become more apparent.

**Table 1.** Human Hematopoietic Stem and Progenitor Definitions.

| HSPC Cell type | Abbreviation | Lineage Potentials | Immunophenotypes |
| --- | --- | --- | --- |
| Hematopoietic Stem Cell | **HSC** | All blood-cell lineages, with self-renewal | Lin- CD34+ CD38- CD90+ CD45RA- |
| Multipotent Progenitor | **MPP** | All blood-cell lineages, no self-renewal | Lin- CD34+ CD38- CD90- CD45RA- |
| Lymphoid-Primed Multipotent Progenitor | **LMPP** | Mo, Gr, DCs, T, B, NK, | Lin- CD34+ CD38- CD90- CD45RA+ |
| Common Myeloid Progenitor | **CMP** | Ery, Mk, Mo, Gr, DCs | Lin- CD34+ CD38+ CD10- CD123med CD45RA- |
| Common Lymphoid Progenitor | **CLP** | T, B, NK, pDC | Lin- CD34+ CD38+ CD10+ |
| Megakaryocyte-Erythrocyte Progenitor | **MEP** | Ery, Mk | Lin- CD34+ CD38+ CD10- CD123- CD45RA- |
| Granulocyte-Monocyte Progenitor | **GMP** | Mo, Gr, DCs | Lin- CD34+ CD38+ CD10- CD123med CD45RA+ |
| Plasmacytoid Dendritic Cell Progenitor | **pDC** | pDC | Lin- CD34+ CD38+ CD10- CD123+ CD45RA+ |

**Abbreviations**
Lin: Lineage Markers, Mo: Monocytes, Gr: Granulocytes, DC: Dendritic Cells, pDC: Plasmacytoid Dendritic Cells, Ery: Erythrocytes, Mk: Megakaryocytes

In human hematopoiesis, conflicting observations in the lymphoid development prompt a deeper examination of lymphoid potentials. Under the hierarchical model, lymphoid priming begins in lymphoid-primed multipotent progenitors (LMPPs), also known as multi-lymphoid progenitors (MLPs), ^7,8^ and continues in the common lymphoid progenitors (CLPs) ^9^. LMPPs exist in the most immature CD38lo compartment and have lymphoid and myeloid potentials but no erythro-megakaryocytic potentials ^7,8^. Conceptually, CLPs are downstream of LMPPs and can generate all lymphoid lineages (T, NK, B, and pDC), but none of the other lineages ^9^. A recent single-cell transcriptomic study has reported that the CD38lo HSPCs exhibit largely unstable clustering results and are highly interconnected in the nearest neighbor graph ^1^. From their findings, Velten et al suggested that the CD38lo progenitors represent very early transitory states, in which the lineage priming is beginning, rather than discrete cell types. On the other hand, the upregulation of CD10 has been associated with B cell differentiation bias and a high degree of IgH DJ rearrangement ^10,11^, suggesting CD10+ CLPs to be B-lineage-primed progenitors. Thus, current definitions for human HSPCs only identify the very beginning of lymphoid priming and primarily B-lineage committed cells, leaving the intermediate stages of T and NK lymphoid development largely undefined.

Lymphoid potential has been reported outside the human LMPP or CLP compartments. Our single-cell mass cytometry study identified lymphoid-phenotype progenitors with expressions of terminal deoxynucleotidyl transferase (TdT) or IL-7Ra, but not CD10 ^11^. These progenitors were located in between CD38-progenitors and CD10+ cells in the B cell development pseudotime, suggesting a lymphoid-specific gene regulation activated in between LMPPs and CD10+ CLPs. Lin-CD34+ CD38+ CD10-CD45RA+ CD62L^hi^ progenitor cells also exhibited full lymphoid and monocytic potentials ^12^. However, the instability of CD62L in the freeze-thaw cycle ^13^ makes it inadequate for frozen BM samples, which are often used in research settings. Nonetheless, Kohn et al. successfully showed the existence of lymphoid progenitors that are not captured in LMPP or CLP gates. Subsequently, residual lymphoid potential was even reported within the canonical granulocyte-monocyte progenitors (GMPs) ^6^ in the human cord blood. Yet the phenotypic nature of the cells responsible for the lymphoid progenitor activity remained elusive. Altogether, these studies highlight the need for assessment of molecular and functional lymphoid potential that could be linked directly to cellular phenotypes within the current human HSPC rubric.

Considering the frequent conservation of mouse and human hematopoiesis ^14–16^, it is natural to compare the lymphopoiesis of the mice and the humans. Similar to the human CD10+ CLPs, the initially described murine CLPs (Lin-Sca-1^lo^ Kit^lo^ IL-7R+ Thy-1-) ^17^ were criticized for their strong B cell bias ^18,19^. It is now understood that among the LSK IL-7R+ Thy-1-progenitors, Ly6d+ cells are B-cell progenitors, and Ly6d-cells are the progenitors with all lymphoid potentials ^20^. Yet, the corresponding population of Ly6d-CLPs remains ambiguous in humans, emphasizing the knowledge gap in the human lymphopoiesis.

In this study, we took a bottom-up, data-driven approach to identify lymphoid progenitors within the single-cell proteomic landscape of human BM HSPCs. We hypothesized that by simultaneous quantification of cell surface and intracellular proteins we could infer the lymphoid lineage potentials and consolidate conflicting observations in human lymphopoiesis. In doing so, we were able to infer the lymphoid lineage potentials in previously undefined cellular compartments within the human hematopoietic hierarchy. This approach revealed TdT+ hematopoietic progenitors with lymphoid-primed proteomic and epigenetic landscapes within the canonical human GMP immunophenotypic compartment. Prospective isolation of this putative lymphoid-primed progenitor population via CD84^lo^ expression confirmed its robust lymphoid potential in functional differentiation assays that was equivalent to that of human LMPPs isolated in parallel. Thus, our data demonstrate the utility of bottom-up interrogation of human systems to define a population based on multi-omic molecular profiling while identifying a significant new source of human bone marrow lymphoid progenitors within a presumed myeloid-committed compartment.

## Results

### Single-cell proteomic map of human bone marrow HSPCs

To create a single-cell proteomic map of human BM HSPCs, we expanded our highly multiplexed CyTOF mass cytometry screen ^21^, enabling quantification of 353 surface protein molecules and 79 intracellular targets, including transcription factors (TFs), histone modification markers (Baskar et al., in preparation), and metabolic enzymes ^22^ (Table S1). While the total 432 targets were split across 15 different staining panels, all the panels included a conserved set of molecules (Figure S1A). Conserved molecules included seven cell surface proteins (CD10, CD34, CD38, CD45, CD45RA, CD90, CD123) used conventionally to define human HSPC types and 4 additional lineage-associated proteins - CD71, also known as transferrin receptor 1, for erythroid lineage ^23–25^, SATB1 for early lympho-myeloid progenitors 1^,26–28^, TdT for lymphoid lineage ^1,11,23^, and IRF8 for DC and monocyte lineages ^29–31^. In total, we analyzed 556,226 CD34+ HSPCs from 3 different donors (Donor 1, 2, 3) and minimized technical batch effects via two rounds of split and pooling to barcode donor and panel information (Experimental Workflow in Figure S1B).

Using a stringent thresholding strategy to reduce the possibility of false positives, we identified 81 screen antigens that are detected in at least 0.1% of CD34+ HSPCs (Figure 1B, S2A). To infer the co-expression data of the 81 targets, grouped cells across the 15 different panels into micro-clusters using the conserved panel and then meta-clustered these micro-clusters using the median expression of both the conserved and screen antigens (Figure S2B, See Methods for details). As a result, we detected 10 clusters within the CD34+ HSPCs and annotated the clusters with prior knowledge, named here A1 to A10 (Figure 1C, 1D).

**Figure 1.**
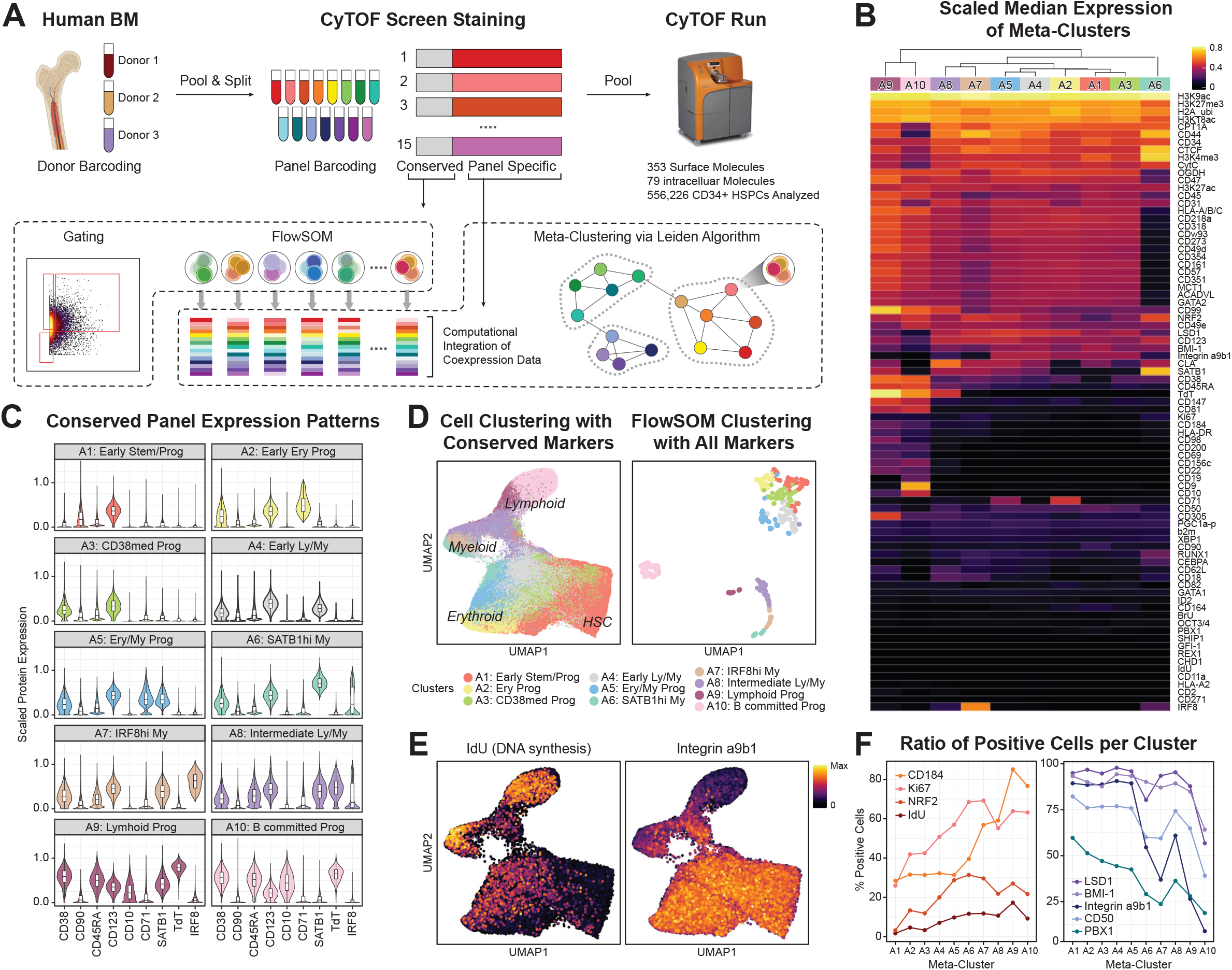
Single-cell proteomic map of human bone marrow HSPCs. (A) Experimental overview. (B) Heatmap of scaled median expression of molecules by meta-clusters. (C) Violin plots of the conserved core panel protein expressions by meta-clusters. (D) UMAP of all CD34+ cells by conserved core panel (left) and of FlowSOM clusters by all protein molecules detected (right). (E) UMAP of all CD34+ cells colored by IdU staining (left) or Integrin a9b1 staining (right). (F) Percentage of positive cells expressing protein molecules that are increasing (left) or decreasing (right) along the hematopoietic differentiation.

Consistent with previous RNA-sequencing (RNA-seq) ^1,23^ and the assay for transposase-accessible chromatin using sequencing (ATAC-seq) ^2^ studies of human HSPCs, we observed CD38lo progenitors to be tightly interconnected with each other, whereas more differentiated CD38hi progenitors were disconnected (Figure 1D, S2B). This observation suggests only subtle differences in the regulatory networks and gene expression of undifferentiated progenitors, consistent with a continuum of low-primed undifferentiated cells (CLOUD) in hematopoiesis ^1^. Among CD38hi progenitors, the CD10+ A10 cluster (B committed progenitors) was embedded distantly from other TdT expressing clusters A8 (Intermediate lymphoid/myeloid progenitors) and A9 (Lymphoid progenitors) (Figure 1C) in the dimensionally reduced Uniform Manifold Approximation and Projection (UMAP) ^32^ space generated with all markers in the panels (Figure 1D, right). This divergence may be due to rapid protein landscape changes as progenitors progress into B cell commitment. Thus, this observation corroborated our hypothesis for the existence of lymphoid progenitors between CD38lo LMPPs and CD10+ progenitors.

Protein-level cell-cycling and epigenetic states of the clusters also correspond to the degree of differentiation. As expected, the most undifferentiated cluster A1 (Early stem/progenitors) had the lowest proliferation marker measurements, including both the lowest 5-Iodo-2′-deoxyuridine (IdU) incorporation (Figure 1E, S2D), which directly labels active DNA synthesis in S phase ^33^, and Ki67 (Figure S2E), whereas more differentiated clusters had higher proliferation (Figure 1F, S2E). H3K27ac histone modification, which marks active enhancers, exhibits a similar pattern (Figure S2E). In contrast, the polycomb complex protein BMI-1 was highest in A1 and decreased gradually in more differentiated clusters (Figure 1F, S2E), as previously reported ^34,35^.

Interestingly, similar trends appeared with several adhesion molecules. Integrin a9b1 heterodimer is an adhesion molecule suggested to play a role in HSPC cell adhesion to endosteal osteoblast ^36^, and exhibited decreased expression level along the differentiation (Figure 1E, S2D). While up to 90% of cells in the earlier clusters (A1∼A5) express Integrin a9b1, this fraction drops to ∼50% of cells in the intermediate A6, A7, and A8 clusters, and to less than 5% in the most differentiated A10 cluster (Figure 1E, 1F, S2D). Similar, if weaker, trends were observed for CD50 and CD164 adhesion molecules (Figure 1F, S2E), emphasizing the link between cell-cell interactions in the stem cell niche to the differentiation processes. Overall, this highly multiplexed single-cell proteomic screen provides deep cellular information on the cell states of human hematopoiesis, via integrating expression patterns of surface molecules, TFs, metabolic states, and epigenetic regulators. Furthermore, the protein molecules highlighted in this screen were used for subsequent analyses and prospective isolation.

### Identification of the proteomic signatures of human BM HSPC populations

While we identified 81 substantially expressed targets from the original screen, we asked if we could identify a smaller subset of these molecules that captured the variance of this larger dataset. To look for redundancy, we calculated the Pearson correlations between all pairs of targets, detecting proteins with highly correlated expression patterns (Figure 2A). Some examples of the most evident correlated proteins observed are lymphoid/B-lineage-associated, including TdT, CD10, CD19, CD9, CD22, and CD200, and stemness-associated proteins, including CD34, CD90, CD164, and PBX1, implying cell-state specific protein expression modules. Given this large amount of structure in the protein expression, we further reasoned that the protein landscape of HSPCs can be recapitulated with a reduced set of representative protein targets from each of these highly correlated groups. To this end, we nominated representative proteins with a differential analysis aimed at capturing the statistically significant differences among protein landscapes of meta-clusters (Figure 2B, Table S2). Putting all of this together, we selected the 23 of the most informative protein molecules from the screen that are either used for conventional HSPCs selection (i.e., gating, Figure 2C, Conventional Gating column) or were identified in our analysis of differentially expressed protein molecules with minimum redundancy (Figure 2C, Top Differential column). Using only these 23 protein molecules, we could recreate original clustering, which used all detected targets from the screen (Figure 2C, right). We finalized our panel with additional lineage markers (Figure 2C, Additional Differential and Lineage column) so as to complete the hematopoiesis trajectories with the more mature, CD34-bone marrow mononuclear cells (BMMCs).

**Figure 2.**
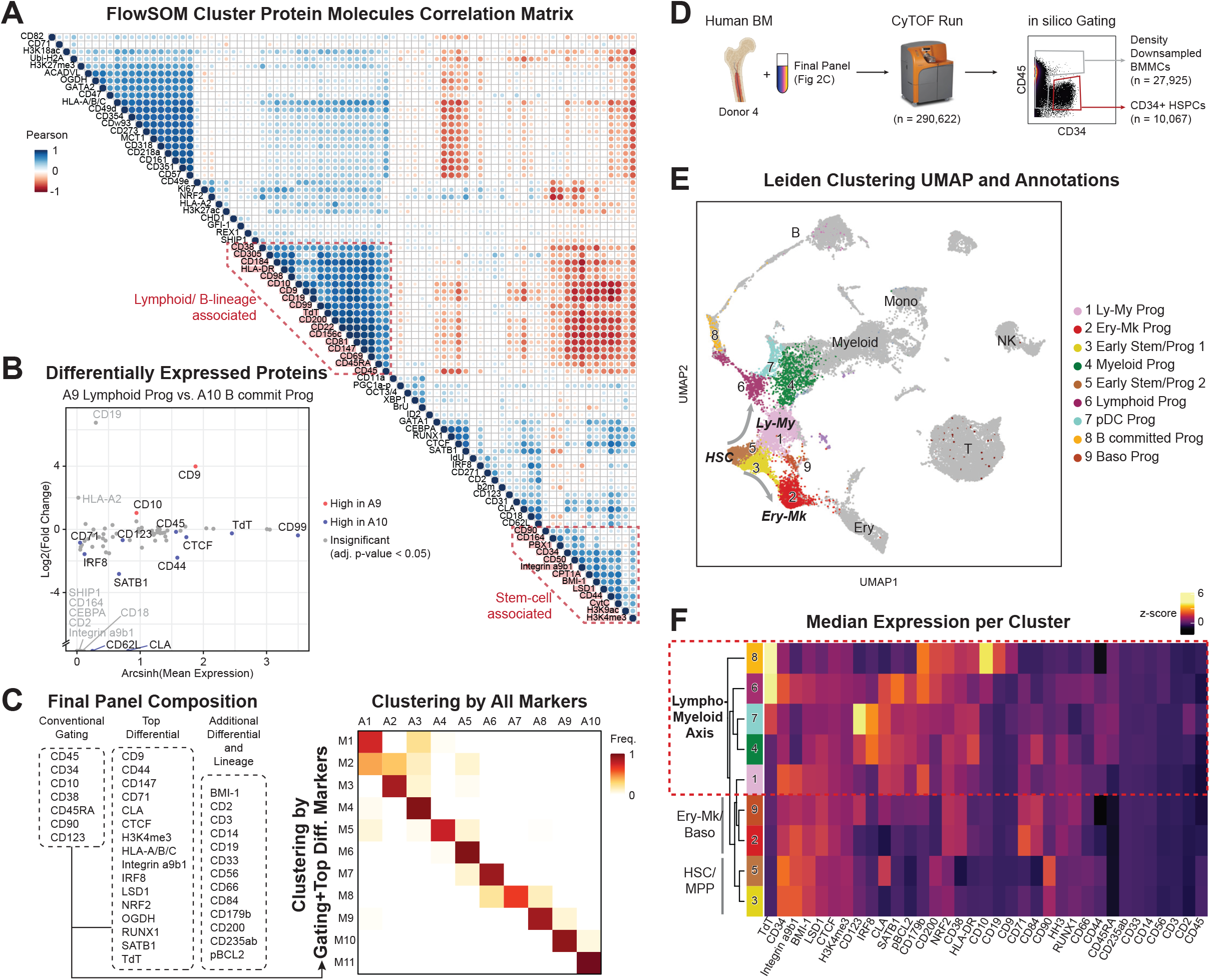
Identification of the proteomic signatures of human BM HSPC populations. (A) Pairwise correlation matrix of protein molecules in the screen in FlowSOM cluster-level (B) Differential analysis of protein expression between A9 Lymphoid progenitors and A10 B-committed progenitors Meta-clusters. (C) Protein targets in the final panel (left) and comparison of meta-clusters from using all markers and minimal set of markers (right). (D) Workflow for the follow-up CyTOF experiment. (E) UMAP of BMMCs. Only CD34+ compartment is colored by their cluster and the rest CD34-compartment is colored gray. (F) Median expression heatmap of protein markers per cluster.

To assess this unified analysis panel, we quantified the protein expressions on a total 290,622 additional cells from another healthy human bone marrow (Donor 4) by mass cytometry. For the emphasis on the hematopoiesis, we enriched HSPCs *in sillico* by gating CD34+ HSPCs and CD34-BMMCs and downsampled the latter population (Workflow in Figure 2D). A nearest-neighbor graph analysis with Leiden clustering demonstrated that the reduced number of single cell proteomic features successfully captured both the heterogeneity of the proteomic landscape among CD34+ HSPCs and the continuum into mature immune cells (Figure 2E, S3A). Within the CD34+ compartment, we were able to annotate 9 clusters (Figure 2E) with their protein expression patterns (Figure 2F, S3B). Of note, we observed bifurcation of Cluster 2 Erythro-Megakaryo (Ery-Mk) Progenitors and Cluster 1 Lympho-Myeloid (Ly-My) Progenitors immediately after exit from Clusters 3 and 5 Early Stem and Progenitors (Figure 2E). Unlike the traditional HSPC model with CMP for erythro-megakaryo and myeloid lineages and CLP for lymphoid lineages, our analysis suggests a shared proteomic phenotype of lymphoid progenitors and myeloid progenitors which separate them from Ery-Mk progenitors. In fact, the Ly-My versus Ery-Mk bifurcation is also observed in transcriptomic and epigenetic landscapes from human HSPC single-cell RNA-seq ^1,23^ and ATAC-seq ^3^ studies, respectively. As the molecular phenotypes in three different modalities coincide, we speculate the functional differentiation potentials to reflect the phenotypic Ly-My versus Ery-Mk bifurcation. Furthermore, we presume the lymphoid progenitors to be sharing phenotypes that have historically been assigned to myeloid progenitors.

### Lymphoid proteomic features identified within the conventional granulocyte-monocyte progenitor compartment

Given the differences between the traditional human hematopoiesis model and our data-driven clustering, we further examined the cluster assignments of canonical HSPC cell types. CD34+ HSPC cell types were annotated based on conventional gating schemes (Figure 3A). We note that our usage of the HSPC cell type names is to refer to the empirical populations with conventional surface immunophenotypes. The most striking observation was found in the granulocyte-monocyte progenitor (GMP, CD34+ CD38+ CD10-CD45RA+ CD123med) compartment, as a substantial subset of GMPs were annotated as Cluster 6 Lymphoid Progenitors (Figure 3B). Cluster 6 was identified as such based on its high expression levels of lymphoid-development-associated proteins, such as TdT and intracellular CD179b, also known as lambda 5 surrogate light chain ^37^ (Figure 3C). In addition, Cluster 6 exhibited lower expression levels of myeloid-associated TF IRF8 and interleukin-3 receptor CD123 in comparison to more apparent myeloid progenitor Cluster 4 and pDC progenitor Cluster 7 (Figure 3C). Moreover, trajectory-inference using PAGA algorithm ^38^ positioned Cluster 6 Lymphoid Progenitor between Cluster 1 Ly-My Progenitors and the Cluster 8 B-committed Progenitors (Figure S4), further corroborating our annotation.

**Figure 3.**
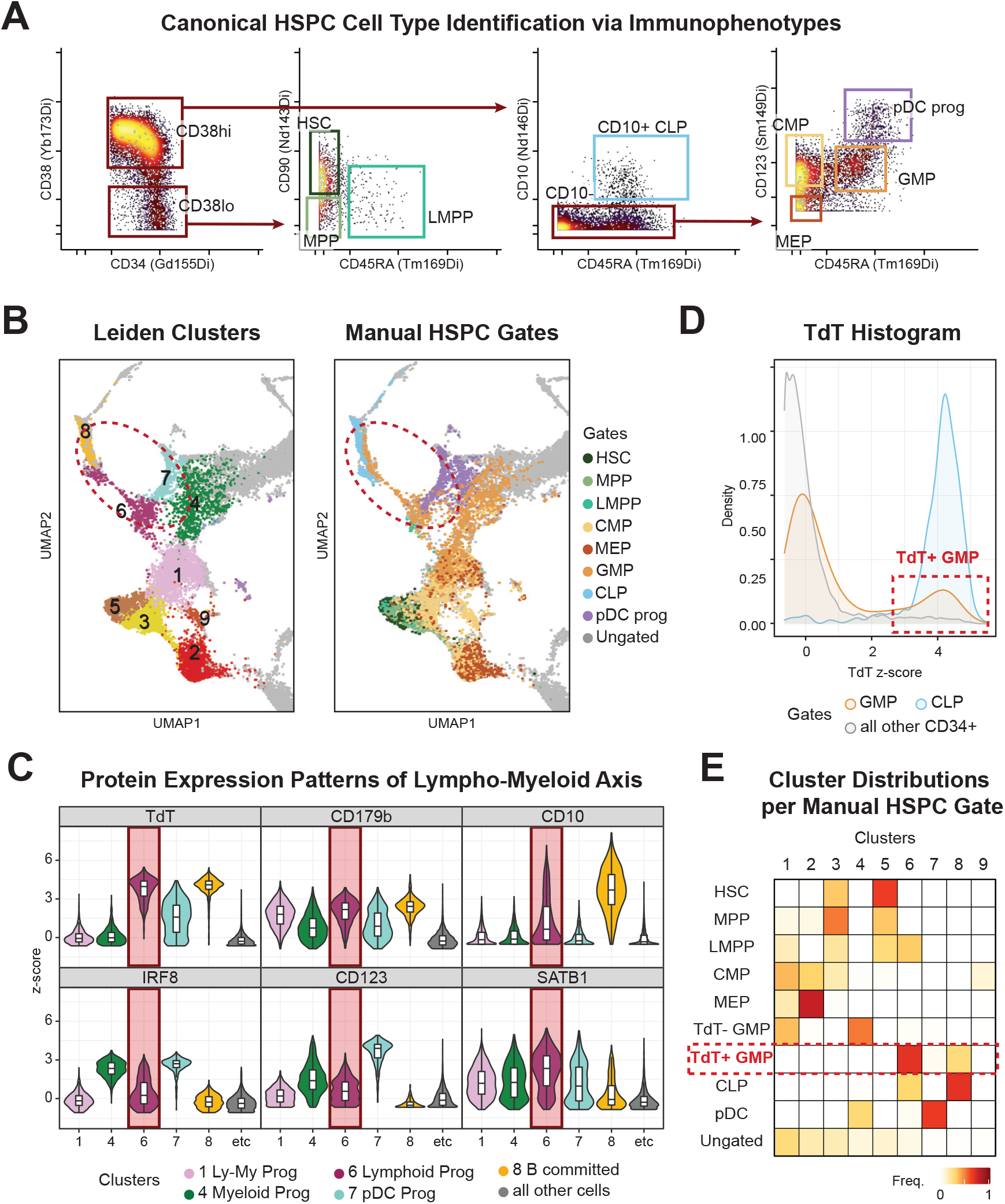
Lymphoid proteomic features identified within the conventional granulocyte-monocyte progenitor compartment. (A) Violin plots of lympho-myeloid lineage associated protein expression patterns. (B) Gating scheme of conventional HSPC cell types on CyTOF. Pregate: Singlet Live Lin-CD45lo CD34+ cells. (C) UMAP from Figure 2E focused on CD34+ compartment. Colored by Leiden clusters (left) or manually gated cell types from Figure 3B (right). (D) TdT protein expression histogram. (E) Confusion matrix of manually gated cell types representing the frequency of Leiden Clusters per cell type.

To delineate the lymphoid progenitors among the cells displaying the conventional GMP surface immunophenotype, we focused on the characteristic expression of TdT of the Cluster 6. TdT, terminal deoxynucleotidyl transferase, is functionally intrinsic to VDJ recombination during lymphoid development and has been identified as the characteristic lymphoid gene in previous studies ^1,11,23^. Furthermore, a recent study in mice demonstrated high TdT expression in lymphoid-biased MPP4s and CLPs but no expression in GMPs ^39^. In contrast, the expression level of TdT in the conventional human GMP compartment was clearly bimodal (Figure 3D) and TdT+ subset had expression level as high as CLPs (Figure S3C). Subsequent classification of TdT+ subset of GMP surface immunophenotype cells (named here TdT+ GMPs) highly corresponded to Cluster 6 based on their proteomic profile (Figure 3E, S3D). Thus, despite its presumed myeloid identity based on the surface immunophenotype, TdT+ GMPs appeared to be previously unappreciated lymphoid progenitors.

### TdT+ subset of human GMPs exhibit a lymphoid-primed chromatin accessibility landscape

Considering the expressions of lymphoid-specific proteins such as TdT, we reasoned that TdT+ GMPs must have already been primed for lymphoid developmental programs. Thus, we assessed the lineage-specific chromatin accessibility of TdT+ GMPs to examine their developmental potentials. Since TdT is an intracellular protein, we utilized inTAC-seq ^40^ to purify target cells based on immunochemistry staining of TdT and assessed the transposase-accessible chromatin from two healthy bone marrow donors (Donor 5, 6). While they both subsets are considered GMPs by their surface immunophenotypes, TdT+ GMPs and TdT-GMPs had 5,915 statistically significant differentially accessible regions between them (Figure 4B). To delineate which transcription factors were associated with these differentially accessible chromatin regions, we applied ChromVAR ^41^ to calculate TF motif enrichment scores. TFs associated with lymphoid development, such as TCF3, TCF4, IRF4, IRF8, and ID4, were highly enriched in TdT+ GMPs (Figure 4C), indicating lymphoid priming of this population. In contrast, multiple members of GATA and CEBP TF families, which are known as canonical myeloid lineage TFs, are strongly enriched in the open chromatin of TdT-GMPs (Figure 4C). Interestingly, we previously identified an enrichment for TCF4 motif accessibility in LMPPs biased towards CLP differentiation, and an enrichment for CEBPE motif accessibility in LMPPs biased towards GMP differentiation ^2^. These same trends were recapitulated in the TDT+ GMPs and TDT-GMPs in our own data.

**Figure 4.**
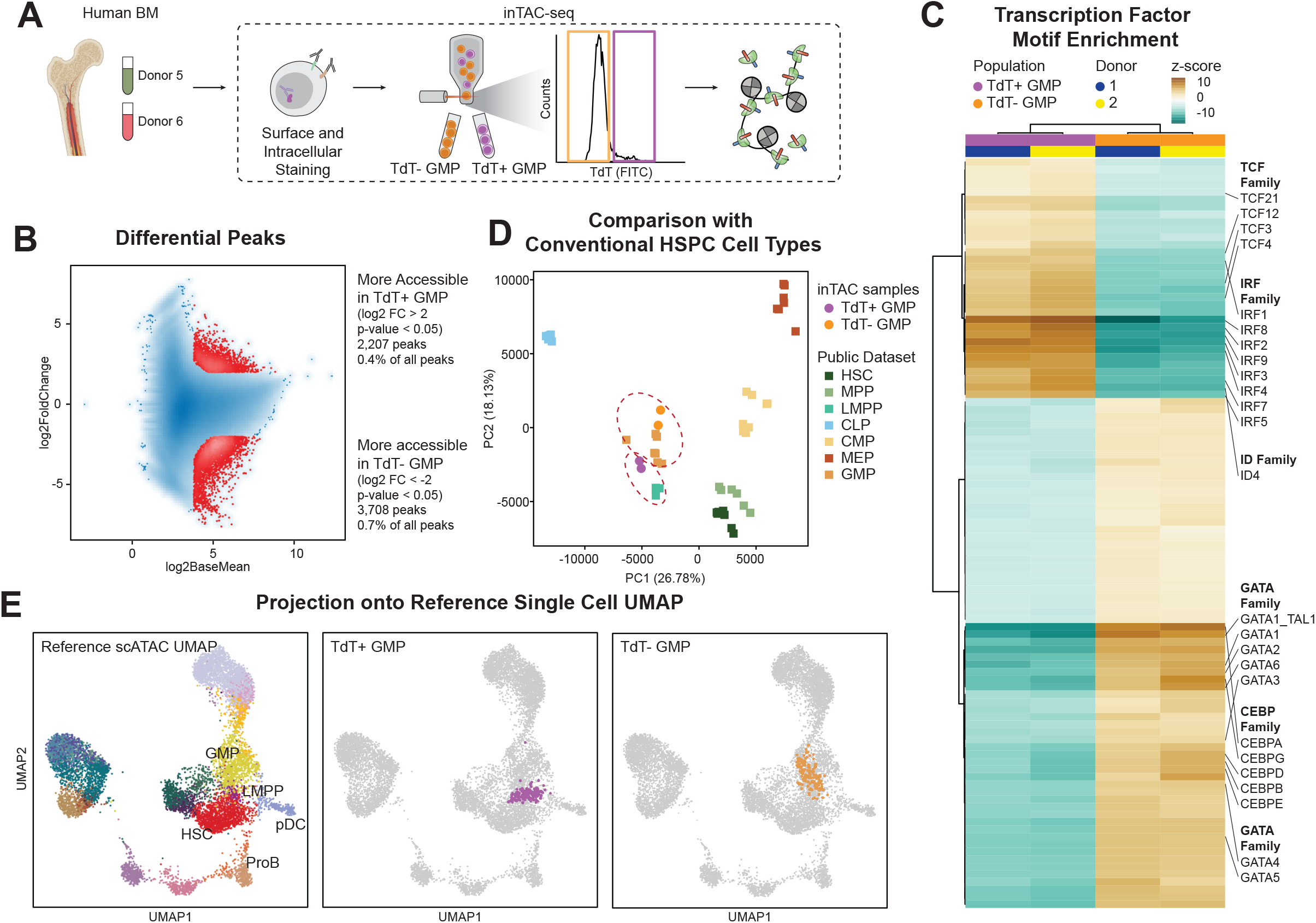
TdT+ subset of human GMPs exhibit a lymphoid-primed chromatin accessibility landscape. (A) Workflow of inTAC-seq. (B) Differential analysis of all peaks detected with TdT+ GMP and TdT-GMPs. (C) Heatmap of ChromVAR transcription factor enrichment scores in differentially accessible peaks between TdT+ GMPs and TdT-GMPs. (D) Principal component analysis of chromatin accessibility from inTAC-seq data from this study with publicly available bulk HSPC ATAC-seq data. (E) UMAP projection of TdT+ GMP (middle) and TdT-GMP (right) inTAC-seq data onto the reference BMMC scATAC-seq data (left).

We then compared the chromatin landscape of TdT+/- GMPs to a bulk human HSPC ATAC-seq dataset ^42^ comprising canonical HSPC cell types identified by surface proteins. When we visualized our data with this data using Principal Component Analysis (PCA), TdT+ GMPs and TdT-GMPs straddled the canonical GMPs from the public dataset, with TdT+ GMPs located more proximal to LMPPs (Figure 4D). This observed grouping and overall proximity of TdT+ GMPs to LMPPs and TdT-GMPs with canonical GMPs was also observed using UMAP visualization (Figure S5). We next compared the chromatin landscape of TdT+ GMPs from inTAC-seq data to a single-cell ATAC-seq data set that spanned the continuum of HSPC differentiation states, independent of the surface-based cell type identification. To achieve this, we simulated single-cell data from our bulk data by subsampling pseudo-single-cell data, then projected these simulated single cells into the original UMAP generated from the reference data (Figure 4E). We observed that TdT+ GMPs overlap with clusters annotated as LMPPs in the direction towards pDCs and Pro-B cells. In contrast, TdT-GMPs overlap with the cluster annotated as GMPs in the reference dataset. Together, the chromatin accessibility of TdT+ and TdT-GMPs suggest that TdT+ GMPs are lymphoid-primed progenitors closer to LMPPs. The myeloid-specific progenitor identity presumed in the surface phenotypic GMPs seems to be restricted to TdT-GMPs.

### Low CD84 surface expression is a surrogate surface phenotype for TdT+ lymphoid-primed progenitors

While TdT is an enzyme with a well-known function in lymphoid development, the measurement of this intracellular protein requires fixation and permeabilization to be detected in primary human cells. Therefore, to prospectively isolate live TdT+ GMPs, we stained an additional aliquot of BMMCs (Donor 4) with a mass cytometry panel focusing on cell surface proteins identified as candidates from our previous mass cytometry experiments or suggested from literatures as lymphoid markers. Then, we computed Pearson correlation coefficients between cell surface proteins and TdT expression levels in the conventional human GMP compartment. (Figure 5A). Among candidates, CD84, also known as signaling lymphocyte activation molecule (SLAM) family member 5 ^43^, had one of the highest absolute levels of correlation (R = -0.429) and a clear bimodal expression pattern in HSPCs (Figure 5B, 5C). Expression of CD84 was previously associated with lineage commitment among CD34+ progenitors ^44^. Other candidates with comparable correlations captured only a small fraction of the TdT+ lymphoid primed population and had a less stark distinction between positive and negative populations (Figure S6A, S6B). Gating on CD84 enriches TdT+ GMPs by 2.3-folds compared to in all GMPs, resulting in 39.2% of CD84^lo^ GMPs expressing TdT (Figure 5D). While our CD84^lo^ GMP gate selected 73.2% of all TdT+ cells among GMPs (Figure S6A), we note that the imperfect selection of TdT+ cells was due to the stringent gating scheme (Figure 5B) and there were less than 2% of TdT+ GMPs in CD84^hi^ GMP gate (Figure 5D). We confirmed that the protein expression patterns of TdT and CD84 were conserved in other donors by additionally analyzing 8 bone marrow samples in a different dataset (Figure S6D) (Favaro and Glass et al., in preparation). Furthermore, CD84^lo^ GMPs and CD84^hi^ GMPs exhibited a clear distinction in their cluster assignments, with CD84^lo^ GMPs corresponding primarily to Cluster 6 lymphoid progenitors and Cluster 1 lympho-myeloid progenitors (Figure 5E). As Cluster 1 lympho-myeloid progenitors preceded Cluster 6 in the trajectory analysis (Figure 3D) and are mostly TdT-negative, we could infer that TdT-CD84^lo^ GMPs were likely earlier progenitors with lymphoid potentials before TdT upregulation. In contrast, CD84^hi^ GMPs were almost exclusively assigned to Cluster 4 myeloid progenitors (Figure 5E). Given these results, we concluded that CD84^lo^ GMPs could represent the lymphoid-primed progenitors in the human lympho-myeloid axis that could be isolated for downstream cellular assays.

**Figure 5.**
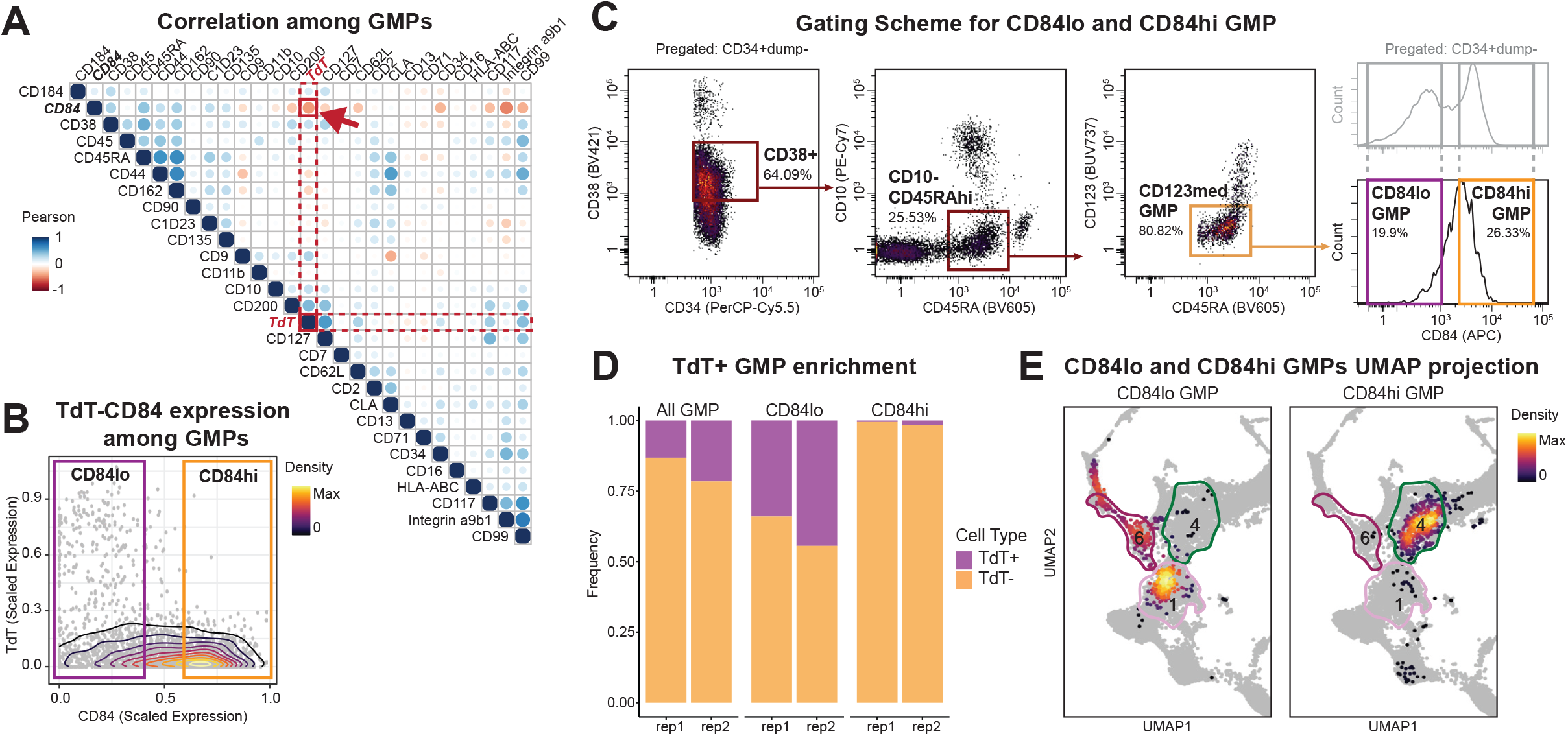
Low CD84 surface expression is a surrogate surface phenotype for TdT+ lymphoid-primed progenitors. (A) Pairwise correlation matrix of candidate surface protein molecules and TdT. (B) Biaxial plot of TdT and CD84 protein expressions in conventional GMPs. (C) FACS gating scheme for CD84lo GMPs and CD84hi GMPs. (D) Frequency of TdT+ GMPs and TdT-GMPs in gated populations. (E) Projection of CD84lo GMPs and CD84hi GMPs in the proteomic UMAP from Figure 2E.

### CD84^lo^ GMPs yield robust multi-lymphoid output with in vitro differentiation assays

To functionally validate the lymphoid developmental potentials presumed from the multi-omic molecular characterization of CD84^lo^ GMPs, we conducted *in vitro* differentiation assays with the OP9-DL4 co-culture system, which can support both T and NK lineage differentiation from human HSPCs ^45,46^. We prospectively isolated CD84^lo^ GMPs from two healthy donors bone marrows (Donor 7, 8) along with CD84^hi^ GMPs, LMPPs, and CLPs (Figure S7A). Consistent with our hypothesis, CD84^lo^ GMPs yielded robust T and NK lineage output (Figure 6C, 6E). By week 3, CD84^lo^ GMPs and LMPPs proliferated significantly more than CD84^hi^ GMPs or CLPs (Figure 6B). While CD84^lo^ GMPs, LMPPs and CLPs all gave rise to lymphoid progeny, CD84^hi^ GMPs only differentiated into CD14+ or CD15+ myeloid cells (Figure 6C). By week 5, CD84^lo^ GMPs and LMPPs continued to expand, but CD84^hi^ GMPs and CLPs yielded few progeny cells (Figure 6D, S7B). CD84^lo^ GMPs and LMPPs had the similar potential for the generation of T lineage cells expressing thymocyte markers, CD1a and CD7 (Figure 6E), and T cell markers, CD4 and CD8 (Figure 6F).

**Figure 6.**
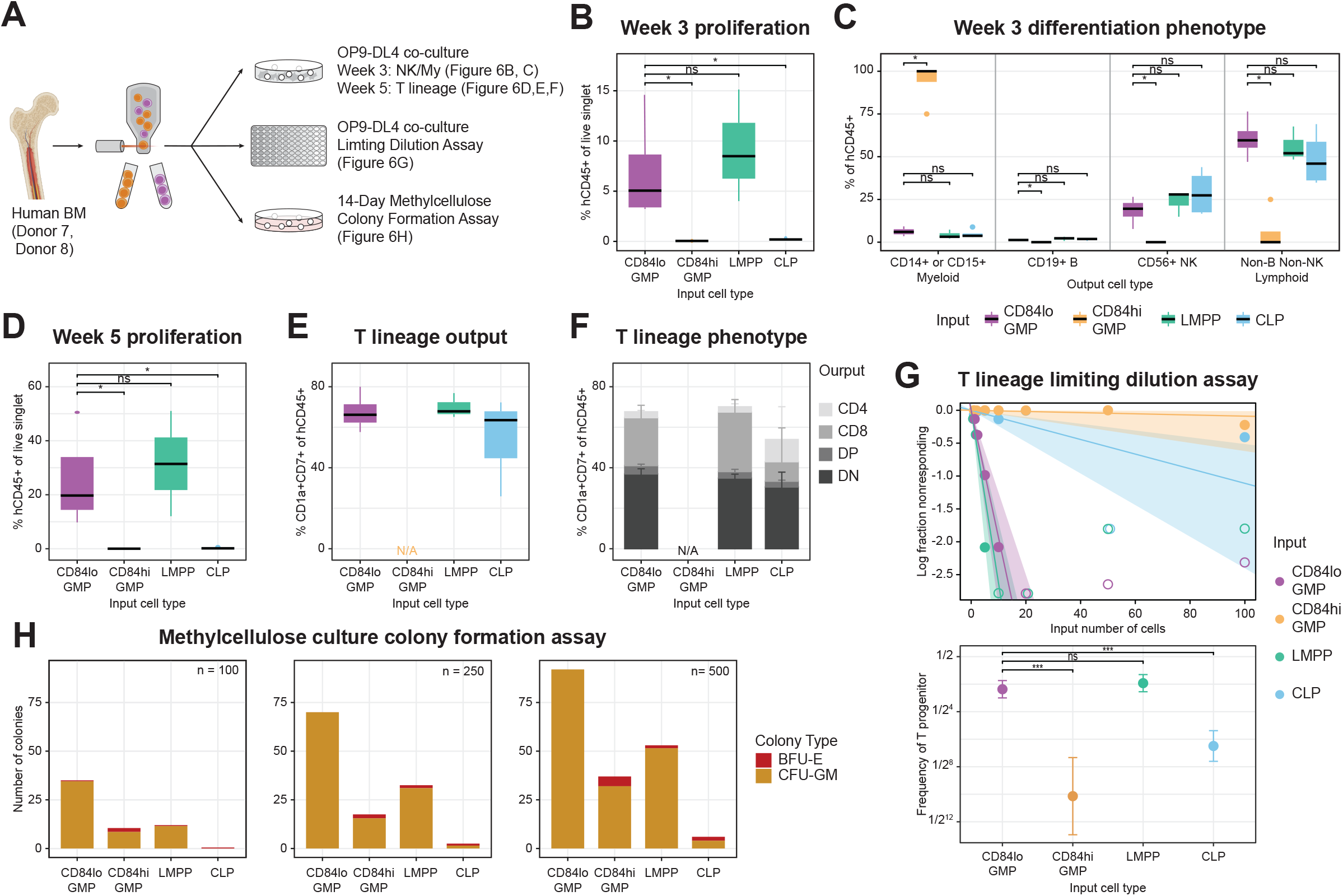
CD84^lo^ GMPs yield robust multi-lymphoid output with in vitro differentiation assays. (A) Workflow of differentiation assays. (B) Box plot of proliferation from OP9-DL4 bulk co-culture in week 3. (C) Box plot of proliferation from OP9-DL4 bulk co-culture in week 5. (D) Box plots of differentiation phenotypes from OP9-DL4 bulk co-culture in week 3. (E) Box plot of T lineage differentiation output from OP9-DL4 bulk co-culture in week 5. (F) Bar plot of T lineage progeny phenotypes from OP9-DL4 bulk co-culture in week 5. (G) Ratio of T lineage progenitor from OP9-DL4 co-culture limiting dilution assay as a line graph of non-responding wells (left) and a summary dot plot (right). (H) Bar plots of methylcellulose colony forming assay with input number of 100 (left), 250 (middle) and 500 (right) cells from each population.

To further quantify the clonal nature and the potency of T lineage potential in these cells, we conducted a limiting dilution assay in the OP9-DL4 system and fitted a generalized linear model GLM using ELDA software ^47^. The estimated frequency (f) of T cell progenitor in CD84^lo^ GMP population was 0.194, similar to that of LMPP population (f = 0.263, pairwise test p-value = 0.345), but significantly higher than CD84^hi^ GMP (f = 8.87e-4, p-value 1.54e-25) and conventional CD10^+^ CLPs (f = 1.11e-2, p-value 1.09e-13) (Figure 6G). Overall, CD84^lo^ GMPs exhibited strong T lineage potential comparable to that of LMPPs, while CD84^hi^ GMPs almost completely lack lymphoid potentials.

In addition, we assessed the erythro-myeloid potentials of CD84^lo^ GMPs via colony formation assay with methylcellulose medium that supports erythroid (E), monocytic (M), and granulocytic (G) lineages differentiation. Consistent with their embedding within the single-cell proteomic landscape (Figure 5E, 3B), CD84^lo^ GMPs consisted of colony-forming unit-granulocytes and macrophages (CFU-GM) cells exclusively, generating approximately 1 GM colony for every 3 input cells (Figure 6H, S7C). This concurrent myeloid potential in CD84^lo^ GMPs reconciles with the conventional “GMP” identity previously associated with these cells. LMPPs or CD84^hi^ GMPs exhibited GM potential to a lesser amount and yielded a few erythroid colonies, suggesting some burst forming unit-erythroid (BFU-E) cells in the input (Figure 6H, S7D). The appearance of erythroid colonies also highlights the promiscuity in lineage priming across phenotypically similar progenitor populations. In contrast, the purity of GM colonies from the CD84^lo^ GMPs highlights the near-optimal purification of lympho-myeloid progenitors by extensive multi-omic molecular characterization. Altogether, our functional differentiation results validate the previously unrealized lymphoid potentials of CD84^lo^ GMPs and demonstrate the prolonged coexistence of lymphoid and myeloid potentials in the human lympho-myeloid axis.

## Discussions

We present a comprehensive summary of the human BM HSPC proteome by quantifying both cell surface proteins and intracellular proteins, encompassing the canonical markers used for cell type identification as well as functional protein molecules that represent the functional state of the cell. Our screen identified 81 protein targets expressed in BM HSPCs and provides a basis on which canonical and modified definitions of HSPC populations can be compared. Using high dimensional proteomic data, we determined the cell types of the clustering results based on the functional proteins. This approach led to the discovery of TdT+ lymphoid progenitors with the traditional immunophenotype of GMPs. Although previously regarded as myeloid committed within the canonical HSPC classification, TdT+ GMPs exhibit lymphoid bias in both proteomic and epigenetic landscapes that are distinct from the rest of TdT-GMPs. Furthermore, we identified CD84^lo^ as a surrogate for TdT+ GMPs for live-cell sorting, where CD84^lo^ GMPs yield robust lymphoid output in cellular differentiation assays. Thus, we report TdT+ GMPs and CD84^lo^ GMPs as previously unappreciated lymphoid progenitors in the human bone marrow.

Our analysis also revealed the erythroid versus lympho-myeloid bifurcation in the proteomic landscape consistent with structures seen in single-cell transcriptomic ^1,23^ and epigenetic landscapes ^3^. Unlike the traditional model with pan-myeloid (including erythroid) versus lymphoid bifurcation in early progenitors, we observe Ery-Mk progenitors (Cluster 2) and Ly-My progenitors (Cluster 1) immediately downstream of the CD38lo early stem/progenitors with multipotency (Cluster 3 and 5; corresponding to HSCs and MPPs in conventional HSPC cell types) (Figure 2E). When we compared conventional gating schemes to the unsupervised clustering of HSPCs (Figure 3E, S3D), we observed cell types in the lympho-myeloid axis, such as CMPs, LMPPs, and GMPs, were dispersed over multiple clusters. This discrepancy between high-dimensional profiling and low-plex cell surface protein-based immunophenotyping in the lympho-myeloid axis emphasizes our current lack of understanding, and possibly over-generalizations, in current models of human early lympho-myeloid development.

In this study, we focused on the protein-level expression of lymphoid signatures, which led to the identification of TdT+ GMPs that correspond to Cluster 6 (Figure 3C, 3D). We noted that despite the distinct lymphoid-primed proteomic and epigenetic landscape, we were unable to define cell surface proteins that separate TdT+ putative lymphoid progenitors exclusively. Instead, we isolated a broader population of CD84^lo^ GMPs. Compared to the previously proposed lymphoid populations, our CD84^lo^ GMPs appeared to be a superset of CD62L^hi^ cells ^12^ and a subset of CD38^med^ GMPs ^6^ (Figure S6C). While the exact differences among these generally overlapping populations should be investigated in future studies, we concluded that the CD84 is the most optimal surrogate for the following reasons. First, the CD84^lo^ GMPs gate most efficiently enriched for TdT+ GMPs (Figure S6A). Second, CD84 expression levels in BM HSPCs exhibited clear bimodal distribution by flow cytometry (Figure 5C). And lastly, TdT-cells among CD84^lo^ GMPs corresponded to the Cluster1 Lympho-myeloid Progenitors, while CD84^hi^ GMPs were identified as myeloid-specific Cluster 4 Myeloid Progenitors (Figure 5E). Thus, selecting for CD84^lo^ GMPs, successfully separates progenitors with lymphoid potential from the myeloid-specific progenitors. Still, it is possible that better prospective surrogates for these TdT+ lymphoid progenitors could be identified in the future. In doing so, we may be able to better assess the level of lympho-myeloid multipotency along the human lympho-myeloid axis.

As B lymphoid cell origins in humans have been well documented ^10,11^, we focused on the T lineage potentials of primary HSPCs in the functional differentiation assays. CD84^lo^ GMPs exhibited robust T cell differentiation *in vitro*, with the frequency of T lineage progenitor of 0.194 from limiting dilution assay. This frequency is ∼17 times higher than that of CD10+ progenitors, highlighting the known B-cell bias of human CLPs ^10^. The existence of CD10-human lymphoid progenitor preceding the B-cell-biased CLPs strongly resembles the identification of Ly6D-all lymphoid progenitors (ALPs) in mice to distinguish from Ly6D+ B-cell-biased progenitor (BLP) ^20^. Considering the evident lymphoid bias in the epigenetic and proteomic phenotype of TdT+ GMPs, we believe that TdT+ GMPs could represent the human equivalent of murine Ly6D-ALPs, and the TdT-CD84^lo^ GMPs are the upstream lympho-myeloid progenitors, the equivalent of human LMPPs or murine MPP4s ^48^.

We recognize that our *in vitro* functional differentiation assays may not precisely represent the *in vivo* potentials of the progenitors. Despite the utility of the xenograft models to study human hematopoiesis, models that support human T lineage development are extremely rare, due to the complicated T cell development process in the thymus. The few existing models, withal, involve complex co-transplantation of human fetal thymic tissues, making them largely inaccessible ^49^. Meanwhile, the *in vitro* differentiation assays we conducted might be more relevant to the clinical application settings. For example, recent studies have demonstrated *ex vivo* differentiation of human T lymphoid progenitors (HTLPs) from BM HSPCs for T cell reconstitution after bone marrow transplantation ^50^ and for T-cell-based immunotherapy ^51^. Thus, the robust *in vitro* T cell differentiation from our lymphoid progenitors in this paper suggest that these cells would likely serve as an effective input for clinical applications.

Our current work does not determine whether the CD84^lo^ GMP population is a lympho-myeloid bi-lineage progenitor population or a heterogeneous mix of uni-lineage lymphoid or myeloid progenitors. However, two recent efforts to link molecular phenotypes to rigorous single-cell functional differentiation assays provide insights. Notta et al. demonstrated a significant reduction of multi-lineage progenitors in BM compared to the CB or fetal liver (FL) ^5^. Karamitros et al. have demonstrated rare lympho-myeloid bi-lineage output in CB HSPCs (only 8% of LMPPs and 7% of GMPs in B/NK and myeloid differentiation condition and 14% of LMPPs and 0% of GMPs in T/B/NK and myeloid differentiation condition) ^6^. Therefore, it is likely that our CD84^lo^ GMP population from adult BM HSPCs is a mixture of phenotypically similar lymphoid or myeloid uni-lineage progenitors with very few to no multi-lineage clones. Nonetheless, we anticipate prospective studies combining lineage tracing and advances in human HSPC differentiation assays to determine the lineage potentials of progenitors *in vitro* and *in vivo*.

While our analysis focused on identifying previously unrecognized lymphoid progenitors, we do not provide evidence supporting strict progenitor-successor relationships among canonical and newly discovered lymphoid progenitors. We trace this challenge to the heterogeneity in canonical HSPC cell types CMPs and LMPPs – the presumed predecessors of GMPs. The heterogeneities among CMPs have also been illustrated both in mice and humans ^5,52,53^.

As we append the lymphoid arm onto this framework, we observe both CMPs and LMPPs dispersed across several of these data-derived clusters (Figure 3E). The top 2 most enriched clusters among conventional CMPs are Cluster 1 lympho-myeloid progenitors and Cluster 2 erythro-megakaryo progenitors, implying the myeloid (Mono-Gran) versus erythroid (Ery-Mk) bifurcation in CMPs. The myeloid progenitors among CMPs are clustered with other lymphoid progenitors (LMPPs, CD84^lo^ GMPs) in Cluster 1 (Figure 3C, S2D), highlighting the similar phenotypes of myeloid and lymphoid progenitors among human HSPCs. Similarly, LMPPs, the other presumed progenitor of GMPs, are scattered across Clusters 1, 3, 5, and 6 spanning the entire lymphoid trajectory prior to CLPs. The dispersed appearance of LMPPs suggests that this surface phenotype marks a molecularly diverse set of cells. Hence, we determined that the canonical immunophenotypes of CMP or LMPP are inappropriate to define predecessor cell types to model progenitor-successor relationships in the lympho-myeloid axis.

Considering the significance of lymphoid development in healthy and malignant hematopoiesis, such as leukemia, bone marrow transplantation, and immune aging, we imagine our characterization of the human bone marrow lympho-myeloid axis to be the basis for future studies and applications. We expect future investigations to query the molecular mechanisms of fate decisions, which will provide the cues to intervene in malignant lymphopoiesis or to boost healthy lymphopoiesis. Beyond the insights to human lymphoid cell potential and cell identity, this study also provides a framework to quantify functional, lineage-associated proteins as surrogates for cell identity within the context of multi-modal single-cell molecular phenotyping. These single-cell molecular archetypes can then be associated with corresponding immunophenotypes for live-cell prospective isolation for functional interrogation and routine enumeration. Via this framework, we successfully reassessed the human lympho-myeloid axis and identified novel, multi-lineage lymphoid progenitors in bone marrow. Moreover, we anticipate our approach of using protein-level molecular regulators of cell function as surrogates for lineage reporters to be expanded to various human tissues beyond the hematopoietic system.

## Methods

### Experimental subject details

Deidentified human bone marrow (BM) (n=8) were obtained from healthy adult donors (AllCells, Alameda, CA).

### Ex vivo labeling human bone marrow for CyTOF screen

Fresh BM aspirates were labeled for their biosynthesis as previously described ^33^. Briefly, Fresh BM aspirates were immediately transferred to a 37°C, 5% CO2 incubator in T75 flask for 30 min prior to SOM3B labeling. A mixture of all three label molecules was added together and mixed thoroughly (final concentration; IdU—100 μM, BRU—2 mM, puromycin—10 μg/mL), and then added to pre-warmed bone marrow. Labeling was conducted in a 37°C, 5% CO2 incubator for 30 min before further processing of BM.

### Human bone marrow processing

Mononuclear cells were isolated from same day BM aspirates by using Ficoll-Paque plus density gradient media (Cytiva) to remove granulocytes and erythrocytes per manufacturer instructions. Bone marrow mononuclear cells (BMMCs) were utilized freshly isolated for the CyTOF proteomics screen and inTAC-seq or frozen in freezing medium (FBS (Omega Scientific) using 10% DMSO (Sigma-Aldrich)) for use in further CyTOF, sorting, and functional assays. Cryopreserved BMMCs were thawed using thawing media (complete RPMI medium [RPMI 1640 (Gibco) supplemented with 10% FBS, Glutamax (Gibco) and 1% Penicillin-Streptomycin (Gibco)], with 20U/mL sodium heparin(Sigma-Aldrich) and 0.025U/mL benzonase (Sigma-Aldrich).

### CD34+ magnetic enrichment

CD34 MicroBead kit (Miltenyi 130-100-453) was used as manufacturer’s instructions to enrich the CD34 compartment from BMMCs in the CyTOF screen and inTAC-seq. CD34-cells in the flow-through from wash steps in the protocol were also washed, counted for cell numbers to be added as a spike-in for the screen or frozen in freezing medium.

### CyTOF antibody preparations

Most CyTOF antibodies were acquired from our previous study ^21^. Additional antibodies needed were conjugated as previously described ^54^ or purchased from Fluidigm. Post-conjugation, each antibody was quality checked on positive and negative control cell lines or human PBMCs and titrated to an optimal staining concentration for 3×1e6 cells per test.

Panel 1∼12 extracellular screening panels were prepared beforehand and lyophilized for storage. Each panel was prepared as a single-test master mix in total 100µL with 100mM D-(+)-Trehalose dehydrate (Sigma-Aldrich) and 0.1X cell staining medium (CSM: PBS with 0.5% BSA and 0.02% sodium azide (all Sigma-Aldrich)) in double-distilled H2O (ddH2O). Prepared single-test aliquots were lyophilized in a vacuum chamber. Lyophilized panels were stored in -20°C until usage. Before staining, each panel was reconstituted in 40µL CSM, pipetted thoroughly, and filtered with Durapore 0.1µm PVDF membrane filter (Millipore) for 2 minutes at 100g to remove any possible precipitates in the antibody mix.

Core panel and Panel 13∼15 intracellular screening panels were prepared on the day of experiment. For all panels, the master mix was filtered with Durapore 0.1µm PVDF membrane filter (Millipore) for 2 minutes at 100g before staining.

### CyTOF screen staining and acquisition (Workflow in Figure S1B)

#### Donor Barcoding and viability staining

CD34+ cells from each donor was supplemented with CD34-cells from the same donor upto total 15 × 1e6 cells in total 100µL with cold FACS benzonase buffer (FBB, FACS buffer [PBS supplemented with 5% FBS and 20U/mL sodium heparin(Sigma-Aldrich)] with 0.025U/mL benzonase (Sigma-Aldrich)) in an 5mL FACS tube. Cells were live-cell barcoded per donor as previously described ^54^. Briefly, each donor sample was labeled with beta-2-microglobulin and CD298 antibodies conjugated with one of In113, Pt195, Pt196 isotopes for 30 minutes at room temperature. Monoisotopic (Pt194) cisplatin (1 μM) was added for the final 5 min to stain non-viable cells ^55^. Cells were washed with FBB and centrifuged for 5 minutes at 300g, 4°C. After removing supernatant by aspiration, cell pellets were resuspended with residual volume and pooled into a single FACS tube and supplemented with CSM upto a total of 300µL.

#### Core panel extracellular staining

Surface staining portion of the core panel was prepared as a 15-tests master mix in total 200µL with CSM. Core panel was added to the pooled sample and stained for 30 minutes. Subsequently, 450µL of FACS Buffer was added to quench staining and the sample was split into 15 FACS tubes with 60µL of sample each, which represent the 15 screening panels.

#### Surface panel staining

Each panel was prepared by reconstituting a lyophilized single-test mastermix as described above. Each tube for surface panels (Panel 1∼12) was added with the 40µL of reconstituted antibody mastermix and stained for 30 minutes. At the end of staining, each tube was washed with PBS and centrifuged for 5 minutes at 250g, 4°C. After removing supernatant by aspiration, cell pellets were resuspended with residual volume.

#### Fixation, permeabilization, and panel barcoding

Foxp3 Fixation/Permeabilization working solution and Permeabilization Buffer were prepared using FoxP3 Transcription Factor Staining Buffer set (eBioscience #00-5523) as manufacturer’s instructions. At the fixation step, 0.5mL of Foxp3 Fixation/Permeabilization working solution was added to each panel tube and vortexed briefly before incubation at room temperature for 1 hour. Cells were washed with 0.5ML of CSM and pelleted by centrifugation for 5 minutes at 600g, 4°C. After removing the supernatant by aspiration, each tube was barcoded with a unique combination of palladium isotopes as previously described ^56^.

#### Intracellular panel staining

Intracellular panel was prepared as a single-test mastermix in total 40µl with Permeabilization Buffer. Cell pellets in each intracellular panel (Panel 13∼15) were normalized to 60µl with Permeabilization Buffer. Each panel mastermix was added to each tube and incubated for 45 minutes at room temperature. After incubation, cells were washed with CSM and centrifuged for 5 minutes at 600g, 4°C. Supernatant was removed by aspiration.

#### Core panel intracellular staining

Intracellular staining portion of the core panel was prepared as a 15-tests mastermix in total 200µL with Permeabilization Buffer. All samples were pooled into a single tube, washed with CSM, and centrifuged for 5 minutes at 600g, 4°C. Supernatant was removed by aspiration and resuspended with Permeabilization Buffer to a total volume of 300µL. Intracellular core panel was added to the pooled sample, briefly vortexed, and incubated for 45 minutes at room temperature. After incubation, the sample was washed with CSM and centrifuged for 5 minutes at 600g, 4°C. After the supernatant was removed by aspiration, sample was briefly vortexed and mixed with 1mL of DNA intercalator solution (1mL of PBS supplemented with 100µL of 16%PFA, 0.5 mM Intercalator-Ir (Fluidigm) and 0.25µM Intercalator-Rh (Fluidigm)). Sample was incubated for 20 minutes at room temperature and then transferred to 4°C for overnight storage before data acquisition.

### CyTOF staining for single panels

Each panel was prepared as two mastermixes, one for surface antibodies and one for intracellular antibodies. Mastermixes were filtered with Durapore 0.1µm PVDF membrane filter (Millipore) for 2 minutes at 100g to remove any possible precipitates. Before staining, BMMCs were suspended in CSM, added 1uL of TruStain FC Blocker (Biolegend) per 1e6 cells, and incubated for 10 minutes at room temperature. Subsequently, cells were stained with surface antibodies in CSM for 30 minutes at RT. All staining volumes were kept to 100µL per 1∼3 × 1e6 cells. After incubation, samples were normalized to a total of 1mL with Low-Barium PBS and added 1µL of 200µM cisplatin (Sigma) for labeling of non-viable cells for 5 minutes. Samples were washed in CSM and fixed using the Foxp3 Fixation/Permeabilization working solution for 30 minutes. Intracellular staining of cells was done by adding antibodies using Permeabilization Buffer for 1 hour. Prior to data acquisition, cells were stained with iridium DNA intercalator solution (1.6% PFA in low barium PBS with 0.5μM Ir191 intercalator (Fluidigm) for 20 minutes at RT or overnight at 4° C

### CyTOF sample acquisition

Before data acquisition, each sample was washed once with CSM and twice with ddH2O. Each wash step was followed by centrifugation for 5 minutes at 600g, 4°C. After the third wash, cells were resuspended with a 1:10 solution of EQ 4 element beads (Fluidigm) in ddH2O to the concentration of 1e6 cells/mL and strained through a 35μM FACS tube filter. Data was acquired on a CyTOF2 instrument (Fluidigm). Single cell events were recorded at a rate of ∼500 cells/second.

### FACS sorting for inTAC-seq

CD34+ enriched samples were washed with PBS and stained with Live/Dead Aqua (Thermo Fisher) as instructed by the manufacturer in dark for 20 minutes. Cells were washed with FBB buffer and spun down at 300g, 5 min, 4°C. Extracellular (E/C) panel (Table S3) was added to samples in FACS buffer in the dark on ice for 30 minutes, followed by a wash with FACS buffer and a spin down at 300g, 5 min, 4°C. For fixation, each sample was fixed with 1ml of 16% PFA for 1 minute before washing with Permeabilization Buffer and centrifugation at 600g, 5 min, 4°C. Intracellular (I/C) panel (Table S3) was prepared in Permeabilization Buffer and added to samples with a brief vortex. Intracellular staining was in the dark on ice for 30 minute.

### FACS sorting for live cells

Thawed BMMCs were washed with FBB buffer and spun down at 250g, 5 min, 4°C. Primary panel (Table S3) was added to samples in FACS buffer in the dark on ice for 30 minutes, followed by a wash with FACS buffer and a spin down at 300g, 5 min, 4°C. Secondary panel (Table S3) was added to samples with a brief vortex, and incubated in the dark on ice for 30 minutes. At the end of incubation, PBS and Live/Dead Aqua were added to the sample and incubated for 20 more minutes in the dark at room temperature. Cells were sorted on a BD FACSAria Fusion (BD Biosciences) at the Stanford Shared FACS Facility.

### inTAC-seq sample processing and library preparation

ATAC-seq samples were prepared as previously described ^40^ with modification. Fixed, permed, sorted samples were spun down at 600g for 5 mins and resuspended in resuspend in 15µl of 1X TD Buffer supplemented with 0.1% NP40. And then added Tn5 in 1X TD Buffer. The amount of Tn5 was normalized to the cell number from the sorter. Cells were incubated at 37°C with 1200 rpm shaking for 30 minutes. 2X reverse crosslinking buffer (2% SDS, 0.2mg/mL proteinase K, and 100mM N,N-Dimethylethylenediamine, pH 6.5 [Sigma Aldrich D158003]) was added at equal volume to transposed cells and reversal of crosslinks was performed at 37°C overnight with 600 rpm shaking. DNA was purified using Qiagen minelute PCR purification columns and ATAC-seq libraries were generated as previously described ^57^.

### OP9-DL4 maintenance

OP9-DL4 cells ^46^ were gifted from Zúñiga-Pflücker lab. Cells were cultured in a 100mm-dish using freshly prepared OP9 media (MEM α, no nucleosides (Gibco12561056) supplemented with 15% FBS and 1% Penicillin-Streptomycin (Gibco)). Cells were split 1:4 or 1:5 when reached 90% confluency as described previously ^45^.

### OP9-DL4 bulk co-culture differentiation assay

1 confluent 100mm OP9-DL4 plate was divided into 2 6-well plates and incubated overnight in a 37°C, 5% CO2 incubator. 24hr after plating OP9-DL4 cells, 600 cells from each sorted HSPC population were deposited into each well and were cultured in the presence of 10□ng/ml SCF, 5□ng/ml FLT3L and 5□ng/ml IL-7 (all from Peprotech, London, UK) in OP9 media (SF7 media). Cells were dissociated from wells by pipetting, filtered with 70µm cell strainer, centrifuged at 300g for 5 minutes and transferred to new plates with fresh OP9-DL4 weekly. Harvested cells were analyzed by flow cytometry at weeks 3 and 5 (Flow panel in Table S3). Wells with fewer than 5 human cells (live hCD45+) were excluded from analysis.

### OP9-DL4 limiting dilution assay

OP9-DL4 cells were plated at a concentration of 2,500 cells in 100µl per well in a flat-bottom tissue culture treated 96-well. 24hr after plating OP9-DL4 cells, different HSPC populations were sorted directly onto 96-well plates. After sorting, 100µl of 2X SF7 media was added in each well. Half of the media was replaced every week. After 2.5 weeks of culture, cells were dissociated by pipetting and transferred to v-bottom 96-well for staining and flow cytometry analysis (Flow panel in Table S3).

### Methylcellulose colony formation assay

MethoCultTM (StemCell Technologies, H4435) was used as the manufacturer’s instruction. Briefly, frozen aliquots of MethoCultTM were thawed overnight. Sorted HSPC populations were diluted with IMDM with 2% FBS to 10X of desired final concentration for seeding. 300µl of diluted cells were mixed to 3ml of MethoCultTM and vortexed thoroughly and then incubated for 5 minutes to reduce bubbles. MethoCultTM mixture containing cells were drawn with a sterile 16-gauge Blunt-End Needle to a sterile 3ml syringe. 1.1ml of the mixture was slowly distributed to a well in 6-well SmartDishTM plates (StemCell Technologies, 27370). 6-well plates were incubated in a 37°C, 5% CO2 incubator for 2 weeks. Differentiation results were counted via STEMvisionTM (StemCell Technologies, 22006) with Color Human BM 14-Day software.

## Quantification and statistical analysis

### CyTOF data preprocessing

Acquired samples were bead normalized using MATLAB based normalization software as previously described ^58^. Sample debarcoding was performed using the premessa R package. Normalized and debarcoded data was then uploaded to either the Cytobank analysis platform (https://www.cytobank.org) or the Cell Engine analysis platform (https://www.cellengine.com). Gated data was downloaded and further analyzed using the R programming language (https://www.r-project.org) and where applicable, Bioconductor (https://www.bioconductor.org) software. Data was transformed using standard inverse hyperbolic sine (asinh) transformation with a co-factor of 5 and column normalized for each individual marker.

### Proteomic screen data integration and clustering

To integrate the data collected on 15 different panels, we first clustered cells into 400 FlowSOM clusters ^59^ (median cell number per cluster: 1288) using the conserved panel. Then each FlowSOM cluster was given the median value for each target in the screen. For meta-clustering, we generated a nearest-neighbor graph of FlowSOM clusters using all median values of 81 targets and then performed Leiden-clustering using FindNeighbors and FindClusters functions, respectively, from Seurat package (https://satijalab.org/seurat).

### Differential protein expression analysis

Differential analysis of protein markers between meta-clusters were performed as previously described ^21^(Glass et al., 2020). Differences in the distribution of molecules were calculated in equally subsampled populations using the KS test. P values were considered significant if the Bonferroni corrected p value < 0.005 to prevent inclusion of false positives in the comparisons.

### ATAC-seq data processing

Adaptor sequence trimming, mapping to the human (hg38 or hg19) reference genome using Bowtie2 and PCR duplicate removal using Picard Tools were performed. hg38 was primarily used for analysis, but the raw data was re-mapped to hg19 genome when comparing to the public dataset mapped to hg19 genome. Mitochondrial reads mapping to “chrM” were removed from downstream analysis. Preprocessed bam files were loaded into R using DsATAC.bam function in the ChrAccR R package (https://greenleaflab.github.io/ChrAccR/index.html). To create a consensus peakset across technical and biological replicates, getPeakSet.snakeATAC function in the ChrAccR package was used.

### Differential accessibility analysis

Differential peaks in the consensus peakset were called by DESeq2 via createReport_differential function in ChrAccR package. Transcription factor motifs enrichment scores in the differential peaks were calculated by ChromVAR package ^41^ via createReport_explanatory function in ChrAccR package.

### inTAC-seq Projection onto scATAC UMAP space

inTAC-seq sample prepared in bulk was projected onto scATAC UMAP space as previously described ^40^. Processed bam files from inTAC-seq samples were downsampled to approximately 150 cells per sample, and loaded into R using DsATAC.bam function in the ChrAccR R package. Count matrix for 500bp tiling regions was converted into a summarizedExperiment data class. Pseudo-single-cells were simulated by subsampling from the summarizedExperiment data class and each pseudo-single-cell was calculated for its UMAP coordinates based on calculating iterativeLSI by using the projectBulkATAC function from ArchR package ^60^.

### Active Progenitor Frequency Estimation from Limiting Dilution Assay

ELDA software (https://bioinf.wehi.edu.au/software/elda/) ^47^ was used for statistical analysis of the limiting dilution assay results.

## Acknowledgments

We thank members of the Bendall and Greenleaf labs for their support and advice. This study was supported by grants from National Institutes of Health 1DP2OD022550-01, 1R01AG056287-01, 1R01AG057915-01, R01AG068279, UH3CA246633 and 1U24CA224309-01 (to S.C.B), RM1-HG007735, UM1-HG009442, UM1-HG009436, 1UM1-HG009442, U2CCA233311, U54-GH010426, and U19-AI057266 (to WJG). This work is also supported by the Defense Advanced Research Project Agency (W911NF1920185 to W.J.G.) and a Stanford Cancer Institute-Goldman Sachs Foundation Cancer Research Award (to W.J.G). WJG is a Chan Zuckerberg investigator. YK is supported by Stanford Immunology Baker Fellowship and by KFAS Overseas PhD Scholarship from Korea Foundation for Advanced Studies. A.A.C. was supported by the NIAID of the National Institutes of Health under award number 5T32AI007290-32 and the National Science Foundation Graduate Research Fellowship Program under grant number DGE-1656518. AT was supported by the Damon Runyon Cancer Research Foundation (DRG-118-16).

## Figure Legends

**Supplementary Figure 1.**
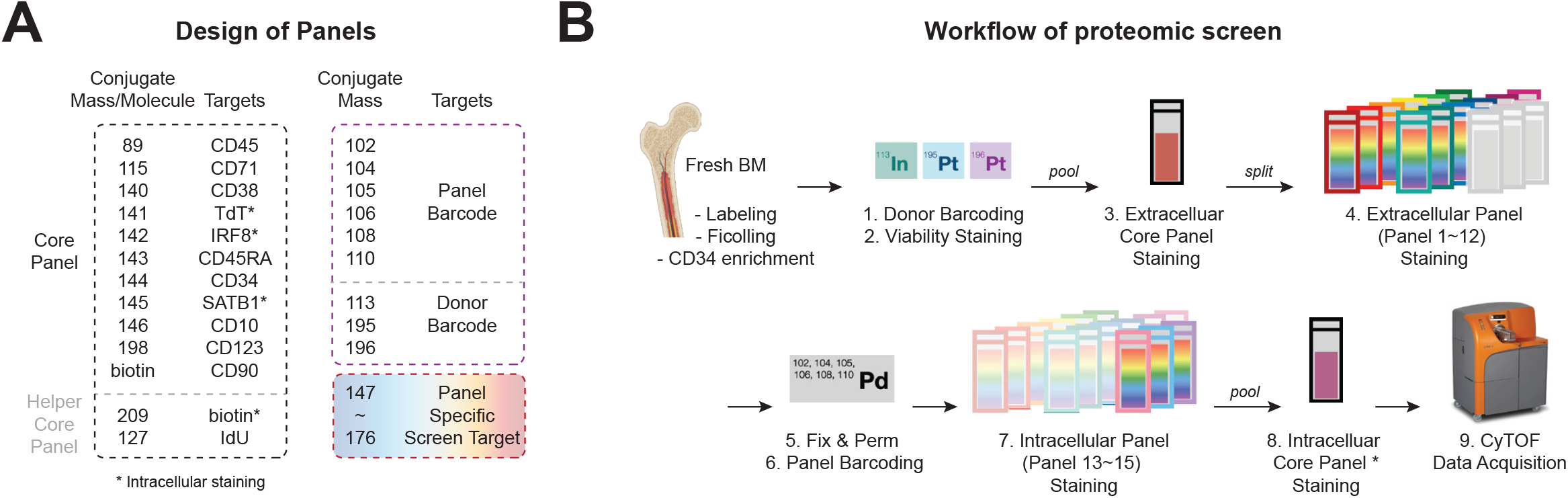
Design of the CyTOF proteomics screen. (A) Design of CyTOF screen panels. (B) Workflow of CyTOF screen.

**Supplementary Figure 2.**
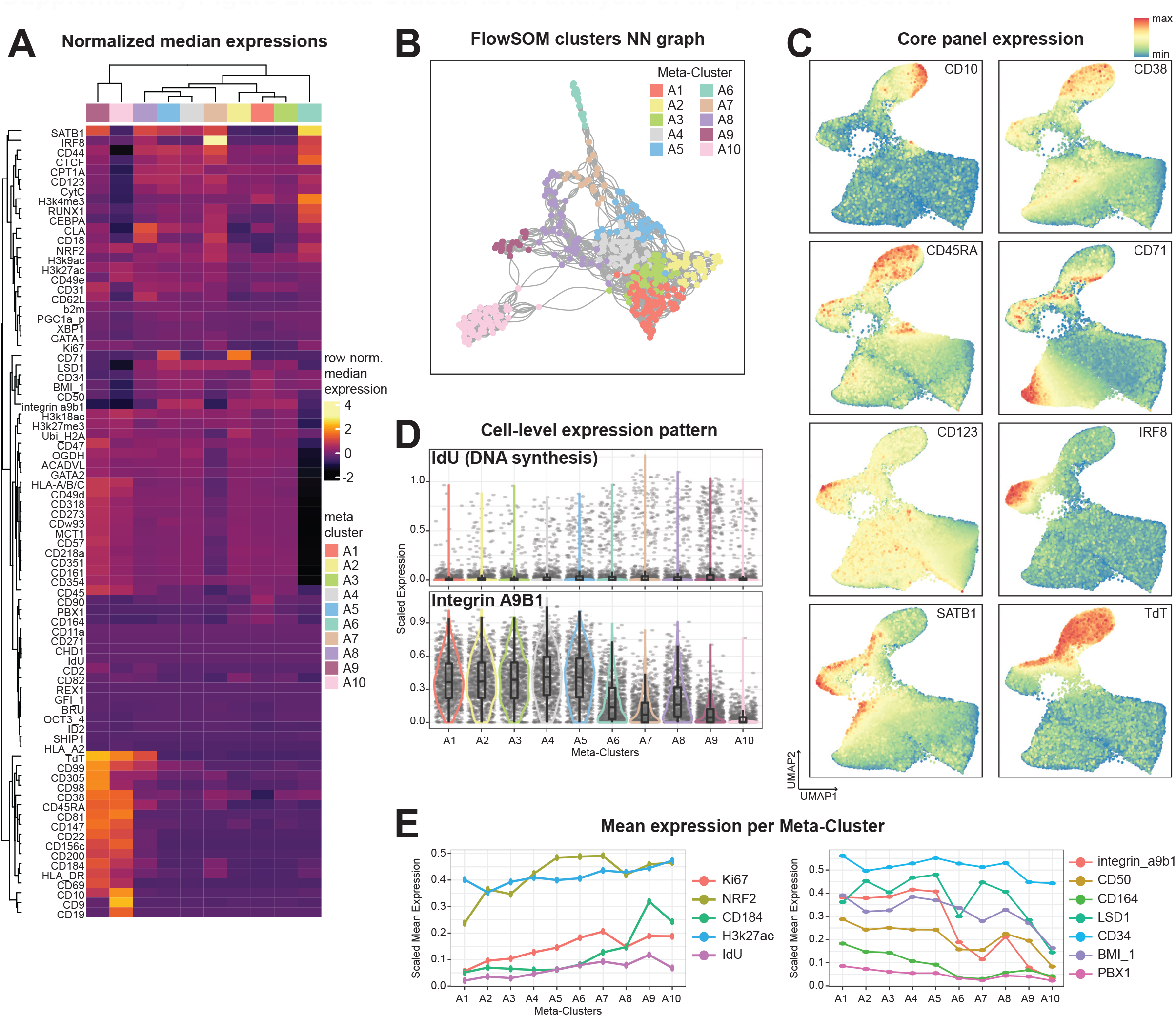
Meta-Cluster level analysis of the proteomic screen. (A) Row-normalized heatmap of scaled median expression of molecules by meta-clusters. (B) Nearest-Neighbor graph of FlowSOM Clusters. (C) UMAP of all cells in CyTOF screen colored by core panel targets. (D) Violin plots of IdU (top) and Integrin A9B1 (bottom). (E) Line graph of mean expression per meta-cluster of protein molecules that are increasing (left) or decreasing (right) along the hematopoietic differentiation.

**Supplementary Figure 3.**
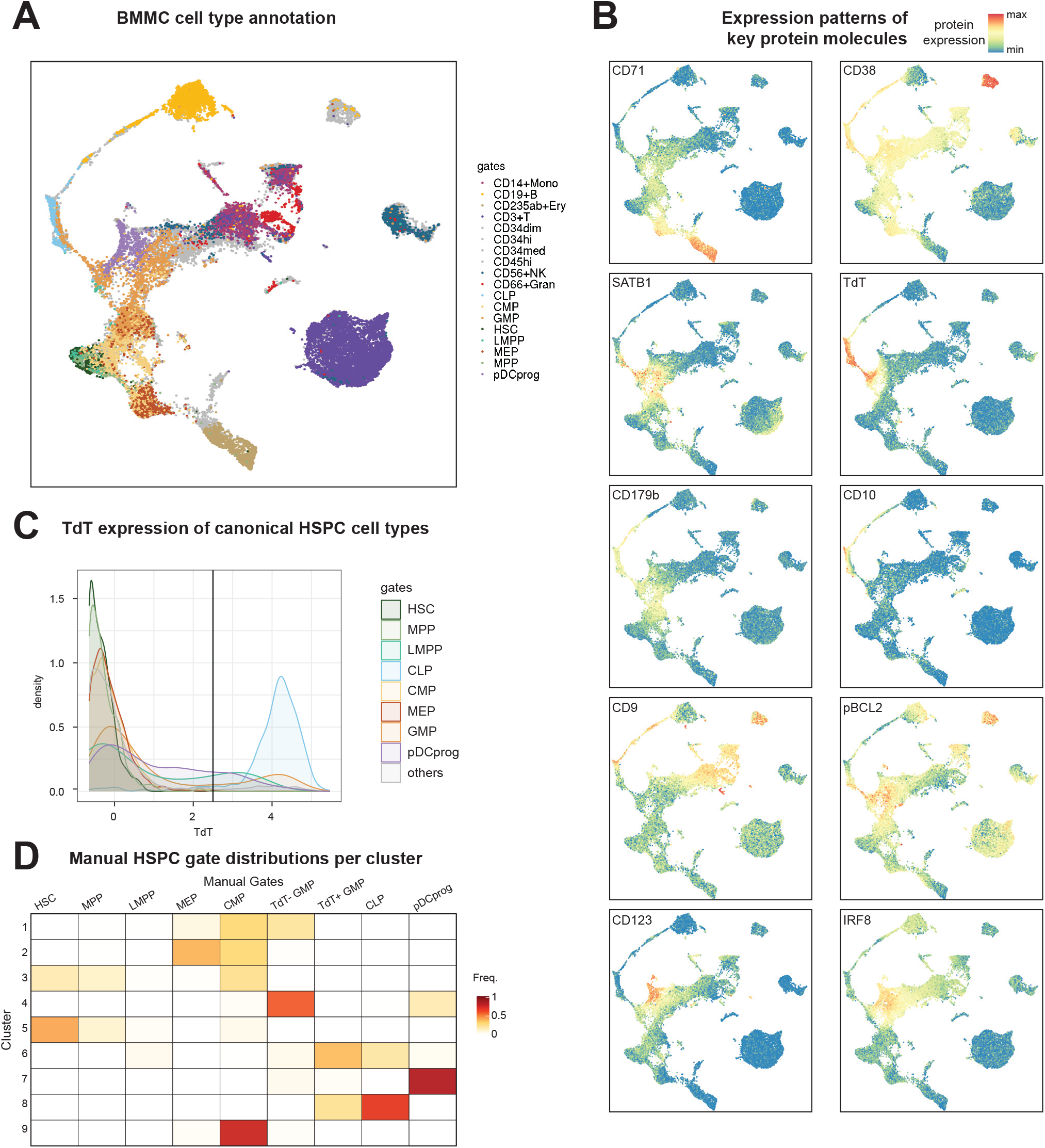
Analysis of BMMC proteome and manual HSPC gates. (A) UMAP of finalized CyTOF panel colored by manual gates. (B) UMAP of finalized CyTOF panel colored by protein molecule expression levels. (C) Histogram of TdT protein expression per HSPC cell type by manual gates. (D) Confusion matrix of Leiden Clusters representing the frequency of manually gated cell types per cluster.

**Supplementary Figure 4.**
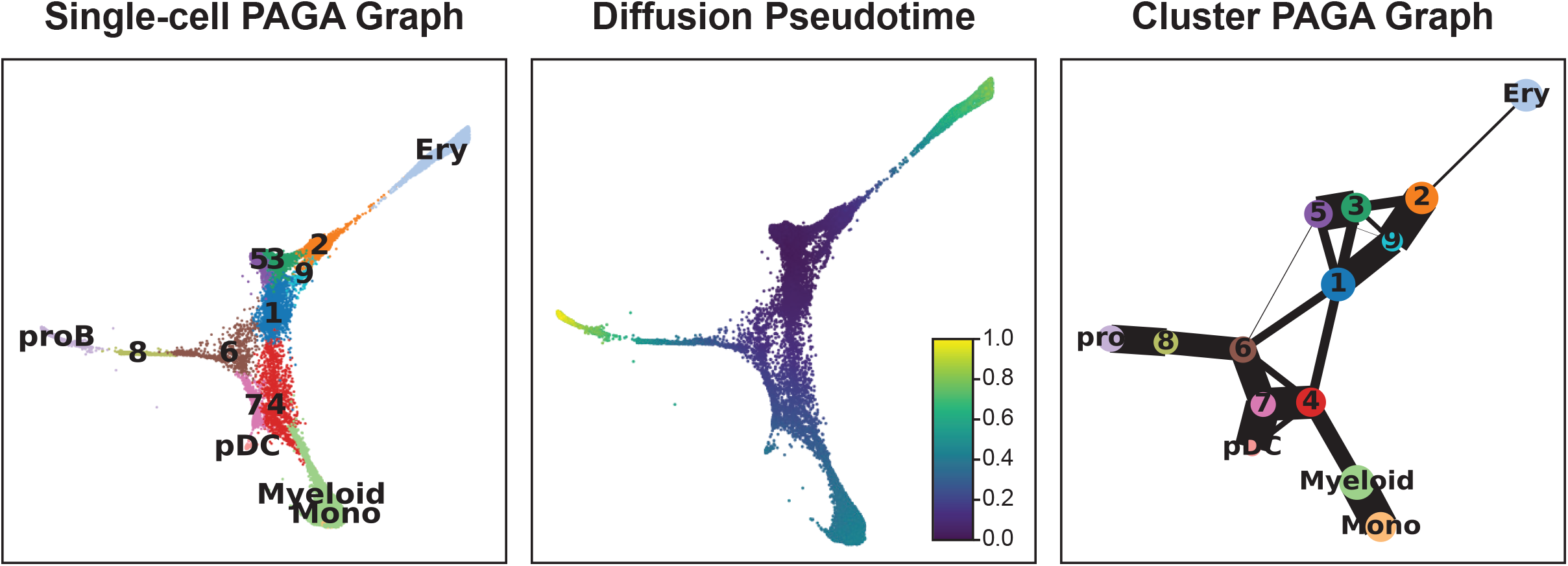
Inference of the developmental trajectory among the HSPC clusters.

**Supplementary Figure 5.**
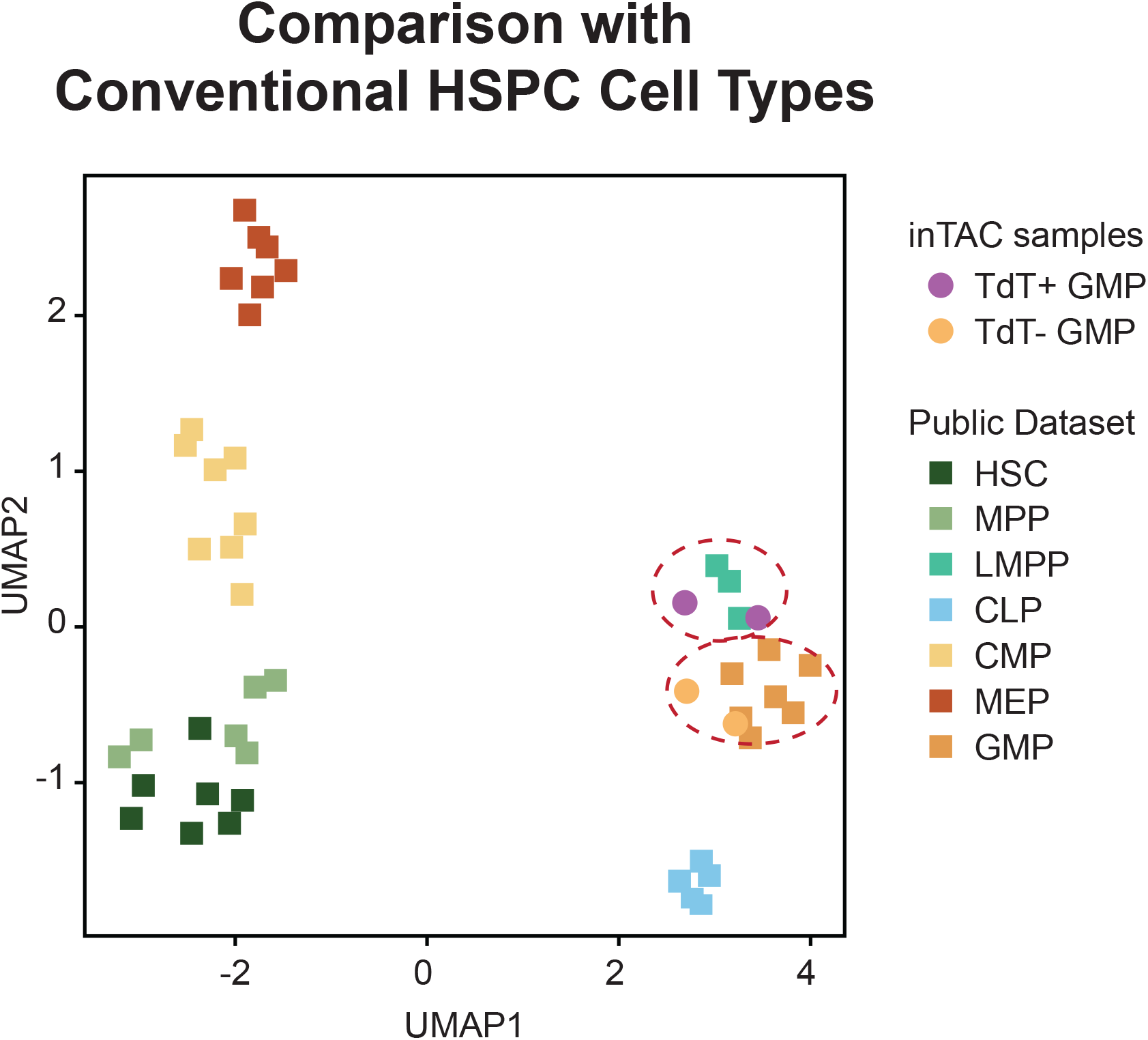
UMAP of chromatin accessibility from inTAC-seq data and control HSPC ATAC-seq data.

**Supplementary Figure 6.**
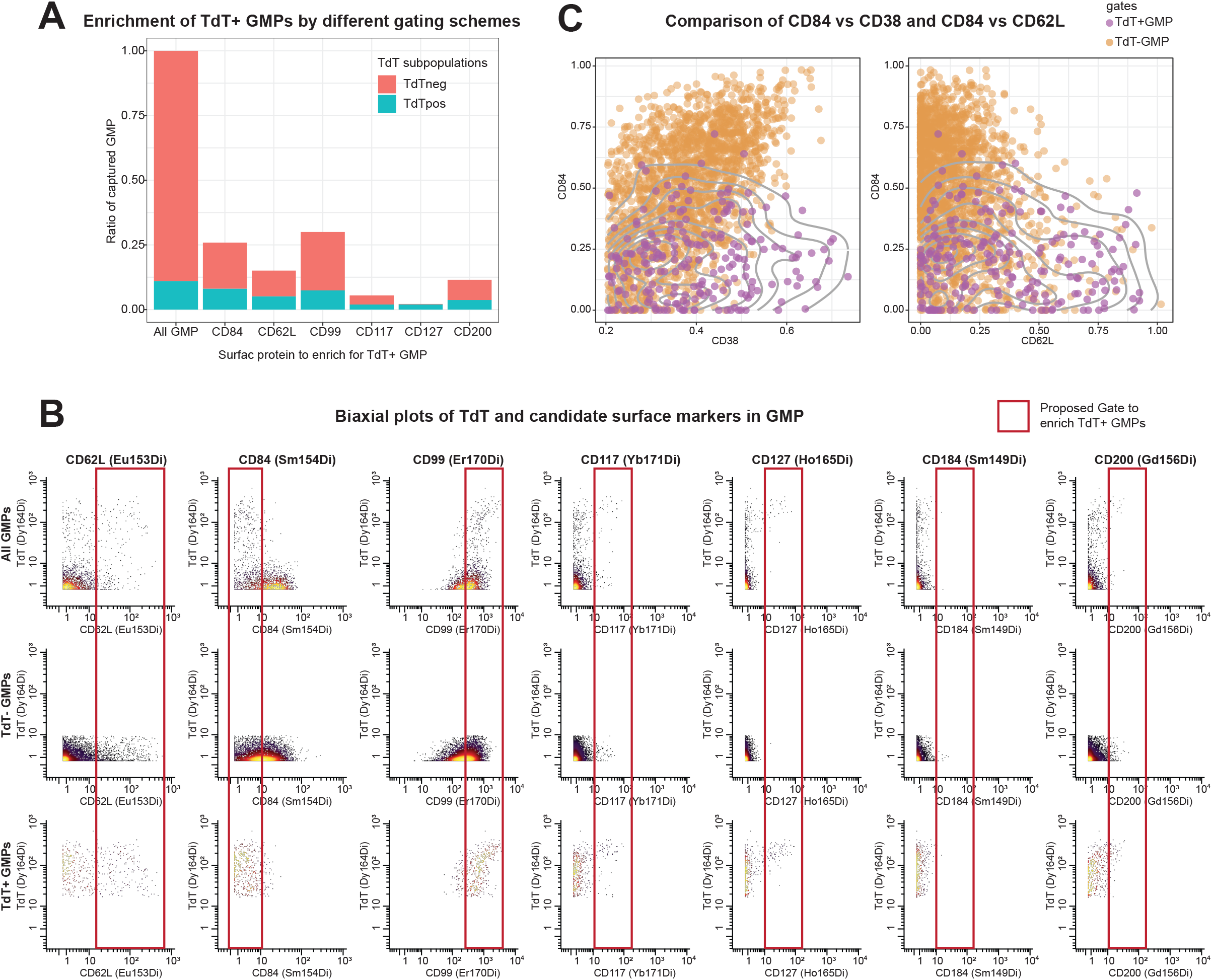
Surface markers to enrich TdT+ GMP. (A) Composition of TdT+ or TdT-GMPs from gating schemes with candidate surface proteins to enrich for TdT+ GMPs. (B) Biaxial plots of GMPs by TdT and candidate surface proteins. (C) Biaxial plots of GMPs by CD84 and previously suggested surface proteins, CD38 (left) and CD62L (right), to distinguish lymphoid progenitors.

**Supplementary Figure 7.**
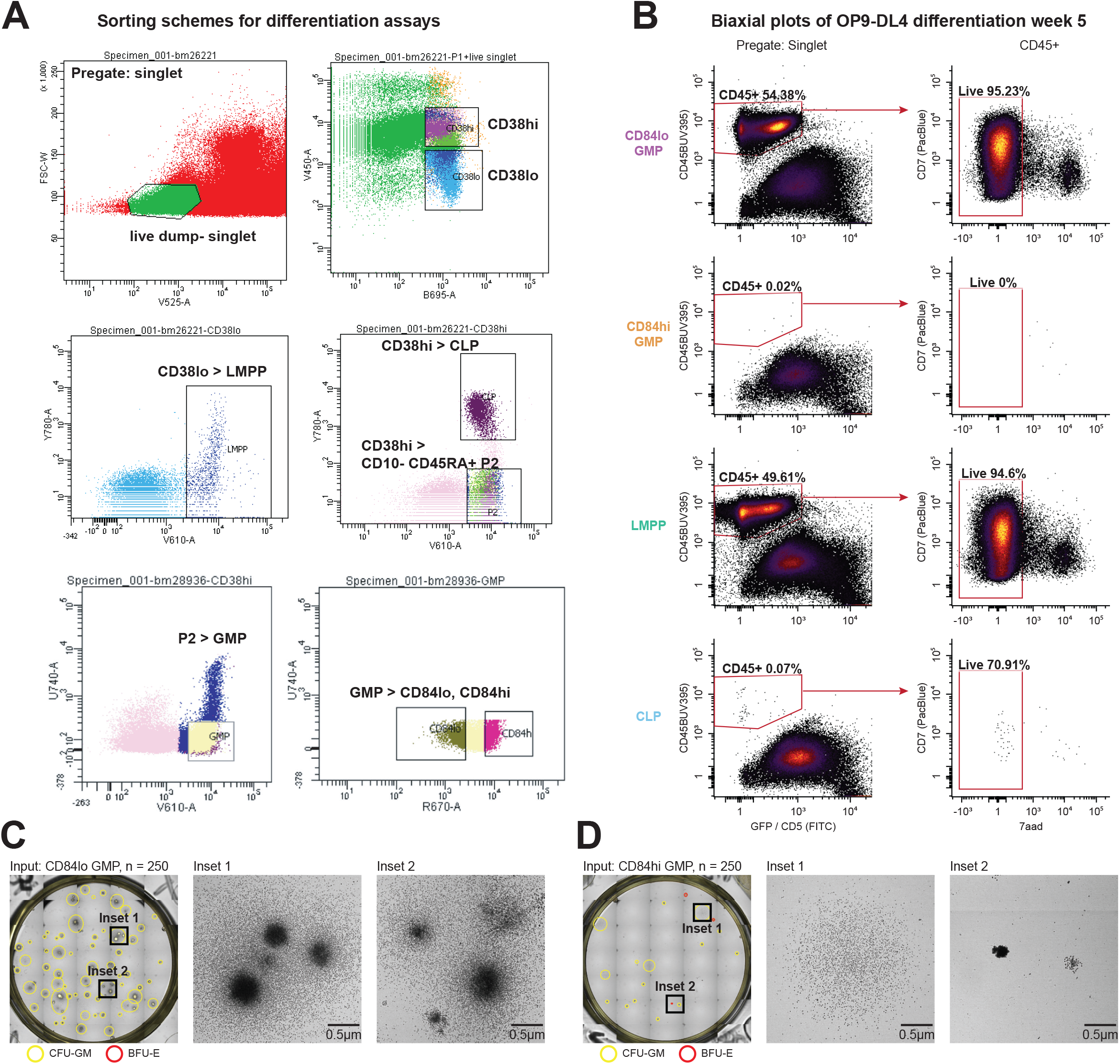
Functional Differentiation Assay Results. (A) Sorting schemes for functional differentiation assays. Representative of the same sorting scheme for all functional differentiation assays. (B) OP9-DL4 bulk co-culture differentiation results after 5 weeks as biaxial plots. (C) Colony forming assay with CD84lo GMPs (n=250) after 2 weeks. (D) Colony forming assay with CD84hi GMPs (n=250) after 2 weeks.

**Supplementary Table 1. Mass Cytometry Screen Panels**

**Supplementary Table 2. MetaCluster Differential Analyses Results**

**Supplementary Table 3. Flow Cytometry Panels**

